# Modeling and targeting haploinsufficiency in *SHINE* syndrome

**DOI:** 10.64898/2026.08.07.743528

**Authors:** Dania Abdellatif, Mustafa Obeid, Rami I. Aqeilan

## Abstract

DLG4-related Synaptopathy, or SHINE syndrome, is a neurodevelopmental disorder caused by *de novo* heterozygous variants in *DLG4* gene, encoding the postsynaptic scaffold PSD-95. Although clinical and genetic evidence support haploinsufficiency, the consequences of pathogenic DLG4 variants in human neurons remain poorly defined. Here, we model three mutations spanning distinct protein domains: a frameshift, nonsense and a missense mutation using iPSC-derived excitatory neurons. Molecular analysis of mature neurons reveals shared PSD-95 deficiency irrespective of transcript levels, together with reduced mature spine density. High-density microelectrode array recordings further reveal convergent and mutationspecific electrophysiological signatures at both single-neuron and network levels, as mutant cultures display genotype-dependent shifts in extracellular waveform states associated with altered firing dynamics. Importantly, restoration of PSD-95 levels using an adeno-associated viral vector (AAV9) harboring human

*DLG4* cDNA and driven by the human neuronal Synapsin I promoter (AAV9-hSynI-DLG4) rescues PSD95 abundance and associated cellular and electrophysiological deficits. Together, these findings establish DLG4 haploinsufficiency as a shared consequence of pathogenic DLG4 variants, while revealing additional variant-associated effects on neuronal structure and activity, rescued by AAV9-mediated neuronal restoration.

## Introduction

DLG4-related synaptopathy, also known as SHINE syndrome, is a newly identified neurodevelopmental disorder associated with variants in the *DLG4* gene, a member of membraneassociated guanylate kinases (MAGUKs). Discs Large Homolog (DLG4) encodes for post-synaptic density 95 protein (PSD-95), the most abundant scaffolding MAGUK protein in the excitatory post synaptic density (PSD)^1^. *De novo* autosomal dominant pathogenic mutations in about 300 patients thus far are associated with moderate to severe intellectual disability, developmental delay, autism spectrum disorder, epilepsy, ataxia, hypotonia, ADHD, language delay, sleep and vision problems^2–5^.

PSD-95 expression is tightly regulated across brain development. In the embryonic mouse brain, PTBP1/PTBP2-dependent alternative splicing of *Dlg4* promotes exon 18 exclusion and nonsense-mediated decay, thereby limiting PSD-95 production. Sequential downregulation of PTBP1 and PTBP2 during neurogenesis permits a gradual increase in PSD-95 expression through postnatal synaptic maturation and into adulthood. Alternative splicing also generates multiple *DLG4* isoforms, with the α-and β-isoforms representing the predominant synaptic forms, the former requiring N-terminal palmitoylation and the latter containing an L27 domain for targeting and stabilization at postsynaptic membranes^6–9^.

PSD-95 is crucial in establishing the complexity of PSDs, through which multiple neurotransmitter systems act, and its role is evident by examining its numerous interaction partners. Indeed, PSD-95 binds and regulates the localization of NMDAR, AMPARs through TARPS, may interact with serotonin and dopamine receptors, in addition to scaffolding proteins like SHANK3 and HOMER1, and is responsible for organization of trans-synaptic cell-adhesion nanocolumns through neurexins (NRXNs) and neuroligins (NLGNs) in pre- and post-synapses, respectively^10,11^. The latter interactions are crucial for synapse formation and maturation, and neurotransmitter receptor recruitment to the membranes^12^. Another fundamental interaction of PSD-95 is with shaker-type voltage-gated potassium channels (KV1) that modulate action potential metrics and resting membrane potential^13^, wherein multiple variants are associated with neurodevelopmental disorders and epilepsy subtypes^14–16^.

Given the diverse signaling pathways associated with PSD-95, patient-identified variants present with disease manifestations consistent with those of its interactome, overlapping with syndromes like PhelanMcDermid, SYNGAP1 and KCNA2^10^. Additionally, in a cohort of 35 epileptic patients examined for developmental epileptic encephalopathy in SHINE patients, moderate to severe ID correlated with epilepsy severity and pharmacoresistance, with age of onset ranging from three months to sixteen years. Epileptiform abnormalities were established to be mainly focal (∼50%) and generalized tonic-clonic (∼40%) seizures^17^.

Considering the nuance of SHINE syndrome and the inaccessibility of human brain tissue for research, and despite increasing clinical and genetic evidence supporting DLG4 haploinsufficiency^18^, the direct molecular, structural and functional consequences of pathogenic variants in human neurons remain undefined. In particular, it is unclear whether distinct variants converge on a shared loss of PSD-95 function, whether they impose additional mutation-specific phenotypes, and whether restoration of PSD-95 is sufficient to rescue the resulting deficits. In this study, we model *de novo* heterozygous DLG4 variants spanning frameshift, nonsense, and missense mutations using iPSC-derived excitatory neurons. We show that these variants converge on reduced PSD-95 abundance and shared synaptic deficits, while retaining convergent and divergent electrophysiological deficits. Importantly, AAV9-hSyn-hDLG4 treatment rescues these deficits, supporting haploinsufficiency as a shared pathogenic mechanism, and AAV9 gene therapy as a viable therapeutic option.

## Results

### SHINE patient mutations converge on a haploinsufficient loss of PSD-95

To gain understanding of the shared molecular and functional signatures underlying DLG4-related Synaptopathy, we focused on three clinically and genetically characterized pathogenic variants associated with severe disease manifestation^2,17^. We employed three different wildtype induced-pluripotent stem cell lines (iPSC) lines for comparison and obtained patient-derived iPSCs harboring a frameshift mutation in exon 7, a nonsense mutation in exon 11 and a missense mutation in exon 19 **(Supplementary Fig. 1A,B**). The latter was modeled using patient-derived (HF) and CRISPR-edited (T654I) lines, see Materials & Methods section. Given the predominant expression of DLG4 in excitatory neurons, we employed the well-characterized NGN2-overexpression system for directed differentiation of iPSCs into excitatory neurons within 21 days of differentiation^19,20^ **(Fig. 1A, Supplementary Fig. 1C**) and evaluated PSD-95 transcript and protein expression in mature (30 DIV) neurons **(Fig. 1B-F, Supplementary Fig. 1D-E)**.

**Figure 1:**
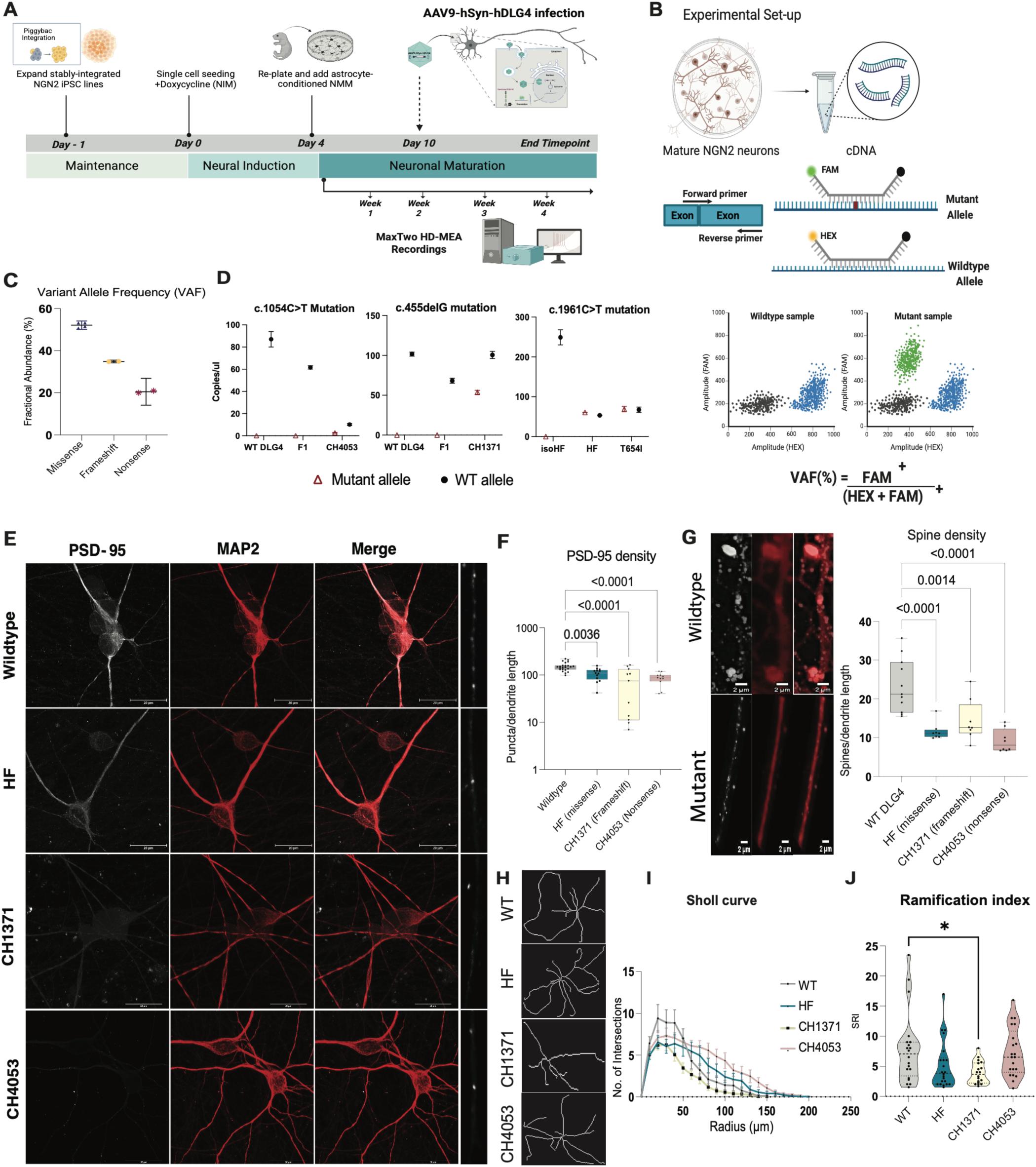
SHINE patient mutations converge on a haploinsufficient loss of PSD-95. **(A)** Workflow schematic of hiPSCs differentiation to NGN2 neurons. **(B)** Schematic of allele-specific expression assay set-up using cDNA of mature 30DIV NGN2 neurons. **(C)** VAF quantification in *DLG4* mutations. **(D)** Mutation-specific quantification of allele abundance in wildtype vs mutant neurons, data points represent individual wells. Wildtype allele in black, mutant allele in red. **(E)** Representative immunocytochemistry for PSD-95 (white), MAP2 (red), across WT and mutant NGN2 neurons at 30 DIV. Scale bars=20µm. **(F)** Quantification of PSD-95 puncta density across 10µm dendritic sections using SynPANal software. **(G)** Representative images of MAP2 labeled, PSD-95 covered spines, with corresponding manual quantification using SynPAnal software. Statistical significance determined through One-way ANOVA, and adjusted for multiple comparisons using Dunnett’s test. Datapoints represent technical and biological replicates across three independent experiments. **(H)** Representative skeletonized tracings of MAP2 labeled neurons**. (I)** Sholl anlaysis of dendritic intersections across 10µm radius intervals from soma center. **(J)** Quantification of Schoenen ramification index across 1µm intervals from soma center using SNT. Statistical significance determined by One-way ANOVA, corrected for multiple comparisons using Dunnett’s test. N= number of neurons from 3 independent experiments. (WT, N= 20), (HF, N=21),(CH1371, N=17), (CH4053, N=18).

First, to determine whether the variants altered transcript abundance, we quantified variant allele frequency (VAF) by droplet digital-PCR (ddPCR) using cDNA from mature wildtype and mutant neurons **(Fig. 1B).** Relative to the expected 50% allele abundance for a heterozygous variant, VAF was reduced for the nonsense and frameshift variants, consistent with degradation of the mutant transcript by nonsensemediated decay in loss-of function variants^21^, whereas VAF for the missense variant remained close to the expected heterozygous ratio **(Fig. 1C-D, Supplementary Fig. 1D)**. To assess the effect at the protein level, we immunostained mature neurons for PSD-95 and the dendritic marker MAP2 and performed immunoblot analysis of 30 DIV mature neurons **(Fig. 1E, Supplementary Fig. 1E).** Quantification of PSD-95 puncta density along 10 µm dendritic segments revealed significantly reduced PSD-95 density in mutants relative to wildtype (missense, p-adj= 0.0028; frameshift, p-adj= 0.0002; nonsense, p-adj< 0.0001) **(Fig. 1F),** suggesting reduced stability of the missense mutant protein.

Next, to discern whether PSD-95 reduction correlated with structural neuronal deficits, we evaluated spine formation in mature neurons, quantifying MAP2 labeled, PSD-95 covered spines across 10 µm dendritic sections^22,23^, and observed significant reduction across all mutants, suggesting destabilization of dendritic spines resulting from loss of PSD-95 **(Fig. 1G)**. Similarly, to evaluate neurite development we performed Sholl analysis, quantifying area under the Sholl curve and the Schoenen ramification index, which represents the ratio of maximum dendritic intersections to primary dendrite number^24^. Overall branching complexity, measured by area under the Sholl curve, did not differ across genotypes **(Fig. 1H,I).** The Schoenen ramification index, however, revealed a mutation-specific phenotype, as only the frameshift mutation showed significantly reduced branching from primary dendrites relative to wildtype **(Fig. 1J).** These results suggest that the reduced PSD-95 content in our patient neurons is translated into structural deficits, consistent with haploinsufficiency of DLG4.

### DLG4 haploinsufficiency in iPSC-derived mutant neurons is associated with convergent and mutation-specific functional deficits

To assess whether the observed molecular and structural deficits are correlated with functional dysfunction, and consistent with epilepsy characterizing severe disease manifestations, we assessed functional neuronal circuit formation using a high-density microelectrode array (HD-MEA) system, comprised of 26,400 electrodes distributed across 1,024 channels^25^ to record extracellular activity. To achieve single-neuron resolution analysis, we performed spike-sorting followed by extraction of singleunit and network-level features using DeePhys, a MATLAB-based platform for functional phenotyping of neuronal cultures^26^.

Firstly, to determine whether DLG4 mutant neurons occupy distinct functional phenotypic space, we performed unsupervised UMAP embedding of DeePhys-inferred features at DIV 21 and observed partially overlapping genotype distributions between wildtype and mutants, and across mutants (WT, *n*= 17 recordings, pooled from three control lines, HF, *n*= 10; T654I, *n*=13; CH1371, *n*=8; CH4053, *n*=12) **(Fig. 2A)**, indicating that genotype-specific phenotypes are not fully resolvable from the full feature space at this stage.

**Figure 2:**
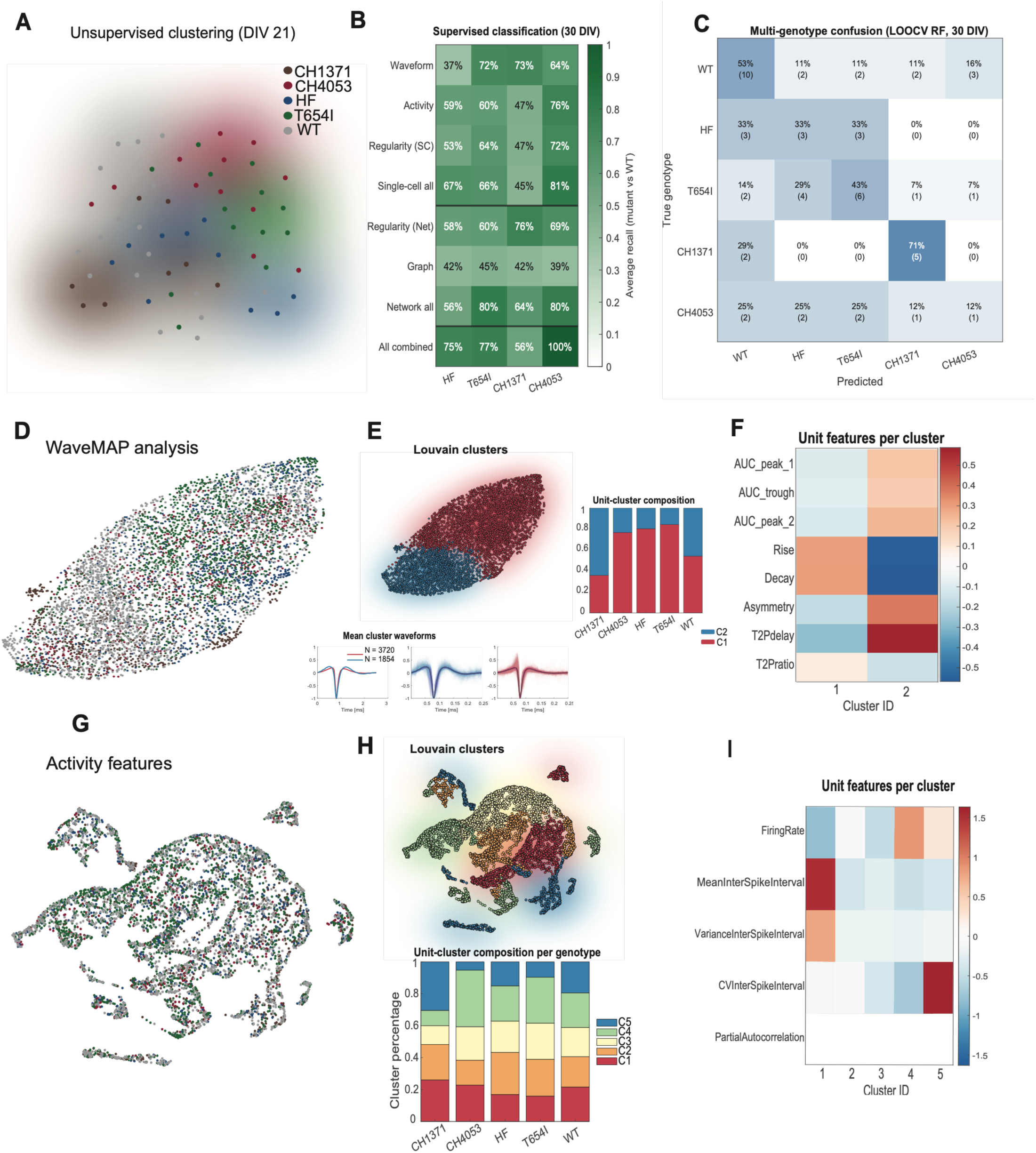
DLG4 haploinsufficiency in iPSC-derived mutant neurons is associated with convergent and mutationspecific functional deficits. KiloSort4 spike sorted HD-MEA recordings of mature iPSC-derived neurons were analyzed in DeePhys across genotypes. **(A)** Unsupervised two-dimensional UMAP embedding at the recording-level colored by genotype and batch normalized. **(B)** Supervised WT versus mutant SVM classification (LOOCV) reporting average recall per feature group and mutant. **(C)** Multiclass genotype confusion matrix (random forest, LOOCV). **(D–F)** Single-cell waveform analysis. (D) UMAP embedding of per-unit waveforms. (E) Louvain waveform clusters with mean waveforms and per-genotype composition. (F) unit-feature enrichment per cluster. **(G–I)** Activity-feature analysis. (G) Per-unit UMAP embedding using activity features. (H) Louvain activity clusters with per-genotype cluster composition. (I) Activity-feature enrichment per cluster. Data collected from 4 independent experiments for untreated cultures, and two batches of treated cultures. *n=* (recordings, units). WT (19, 1827), HF (9, 924), HF AAV9 (9, 589), T654I (14, 1462), T654I AAV9 (7, 953), CH1371 (7, 707), CH1371 AAV9 (8, 572), CH4053 (8, 654), CH4053 AAV9 (7, 339). Total 88 recordings, 8027 units.

To identify which feature classes carry genotype discriminating information, we trained support vector machine classifiers on each mutant line against wildtype at 30 DIV, using single-cell and network feature groups separately and combined **(Fig. 2B),** with samples obtained from 57 recordings and 5,574 units, collected from 3-4 independent experiments (WT, *n*=19 recordings/1,827 units, pooled from three control lines; HF, *n*=9/924; T654I, *n*= 14/1,462; CH1371, *n*=7/707; CH4053, *n*= 8/654). Combining all features yielded high recall among mutants, except for the frameshift mutation (CH1371), consistent with distinct functional phenotypes across variants. Moreover, to discern whether the mutant lines are mutually distinguishable, we performed multi-class random forest classification with leave-one-out crossvalidation **(Fig. 2C).** We observed a higher resemblance of missense mutant lines to each other and to wildtype, whereas the frameshift mutant was recovered with the highest recall, only partially overlapping with the control. On the contrary, CH4053 was misclassified to a similar level across all lines, indicating that the nonsense mutant phenotype, although robustly separated from wildtype in the binary comparison, is not unique among the mutant lines.

To resolve these differences at the unit-level, we applied WaveMAP analysis, embedding extracellular reference waveforms by UMAP followed by Louvain community detection^27^, which resolved two waveform clusters across all units (C1, n = 3,720; C2, n = 1,854) (**Fig. 2D,E**). Cluster assignment was genotype-dependent, whereas wildtype units were distributed approximately evenly between the two clusters, CH1371 units were predominantly assigned to C2, and CH4053, T654I and HF units to C1 **(Fig. 2E)**. Additionally, the clusters were separated principally by repolarization kinetics, with C1 units displaying steeper rise and decay slopes and C2 units displaying prolonged trough-to-peak delay, greater asymmetry, and larger areas under the trough and both peaks **(Fig. 2F),** suggesting a mutation-specific shift in action potential waveform properties.

Finally, to assess whether similar grouping arises from firing dynamics, we performed unsupervised clustering using activity features, which resolved five unit clusters **(Fig. 2G–H).** Cluster composition revealed modest changes with predominant representation of cluster 4 in nonsense and missense mutant lines, and cluster 5 in framshift mutant line, compared to wildtype composition **(Fig. 2H),** with the clusters mainly distinguished by firing rate and interspike interval statistics **(Fig. 2I).** Together, these data demonstrate that the molecular and structural deficits observed in DLG4 mutant neurons are accompanied by measurable convergent and mutation-specific functional alterations at both the single-neuron and network level.

### AAV9-hSynapsin-hDLG4 treatment restores PSD-95 levels and associated synaptic deficits

After observing the effect of reduced PSD-95 abundance on the structural and functional levels in our DLG4 mutant neurons, we aimed to explore the feasibility of using an adeno-associated viral vector serotype 9 (AAV9) to restore PSD-95 protein levels. To this end, we designed and generated two AAV9 viral constructs carrying human DLG4 cDNA for the DLG4 alpha isoform (NM_001321075.3) and beta isoform (NM_001365.5), both under the human SynapsinI promotor to drive specific neuronal expression **(Fig. 3A)**. Firstly, we tested efficiency of neuronal transduction through intracerebroventricular (ICV) injection into wildtype mice at P0-P1, along with a positive control AAV9-hSynI-eGFP virus. We observed robust infection efficiency and biodistribution, most notably in the cortex, as evident by GFP expression **(Supplementary Fig. 2A)** and overexpression of full-form PSD-95 across both isoforms **(Supplementary fig. 2B)**.

**Figure 3:**
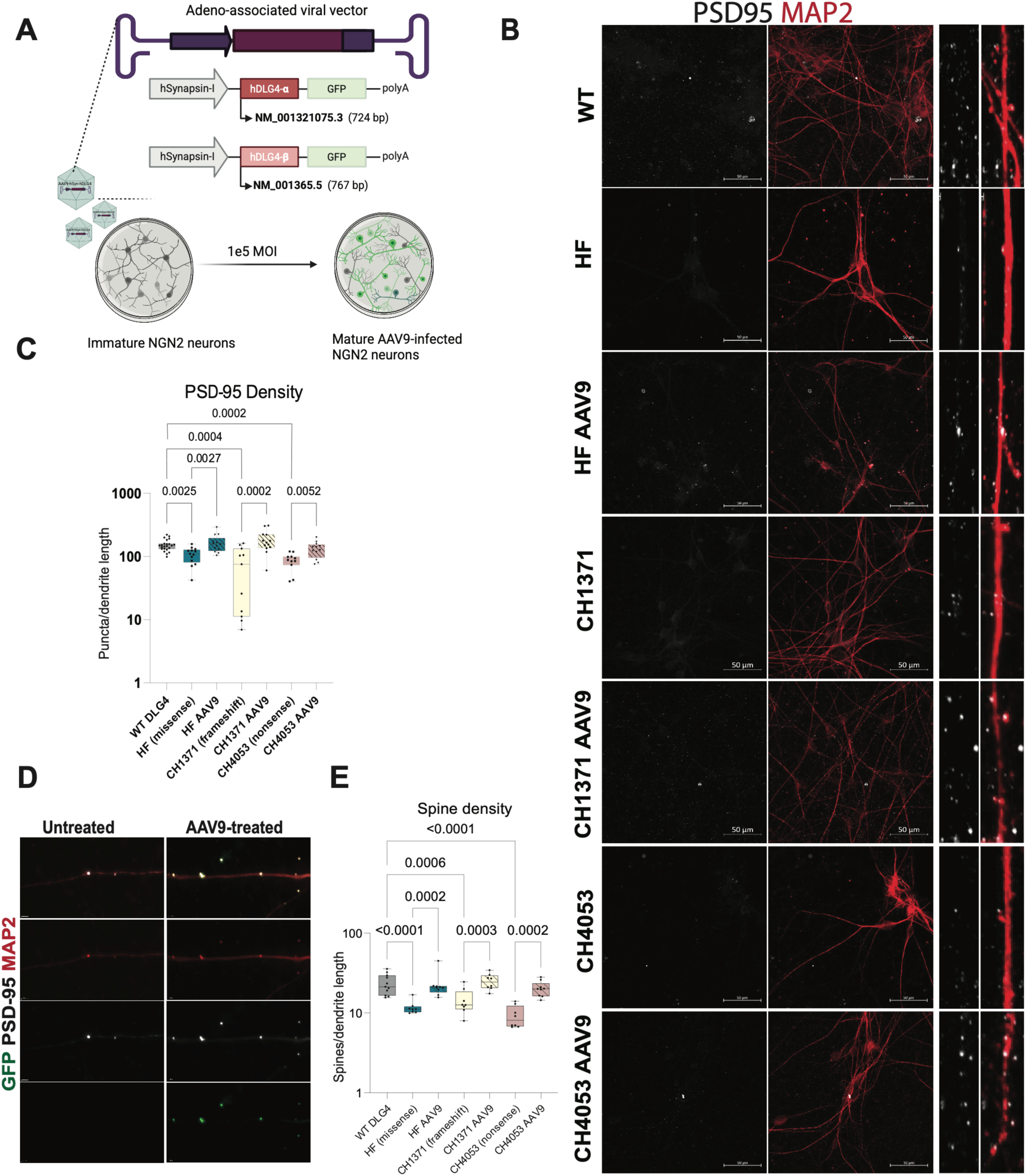
AAV9-hSynapsin-hDLG4 treatment restores PSD-95 levels and associated synaptic deficits. **(A)** Schematic of AAV9-hSynI-hDLG4 α- and β-isoform constructs and NGN2 neuronal transduction. **(B)** Representative images of PSD-95 (white) and MAP2 (red) labeled mature NGN2 neurons and high-magnification of a 10 um dendritic section showing PSD-95 snd MAP2 labeled spine heads. Scale bars= 50 µm. **(C)** Quantification of representative images in (B) showing PSD-95 puncta density. **(D)** Representative images of spines in untreated versus AAV9-hSyn-hDLG4.GFP infected cultures showing GFP colocalization to PSD-95. **(E)** Average spine density across mutant and AAV9-treated cultures. Statistical significance determined through One-way ANOVA, and adjusted for multiple comparisons by controlling the FDR using BKY. Datapoints represent technical and biological replicates across 3 independent experiments.

Given our viral preparation demonstrated efficient neuronal tropism *in-vivo*, we next evaluated its capacity to restore PSD-95 levels *in-vitro*. To this end, we infected immature DLG4 wildtype and mutant NGN2 neurons with AAV9-hSynI-hDLG4 (isoform α), and analyzed cultures at 30 DIV by immunoblot analysis validating expression of full-form functional PSD-95 **(Supplementary Fig. 2C)**. To assess the extent to which AAV9 gene therapy rescues cellular deficits associated with DLG4 haploinsufficiency, we infected neurons with AAV9-hSynI-hDLG4 (isoform α and β), and immunostained against PSD-95 and MAP2 neuronal marker **(Fig. 3B-E)**. Since both vector isoforms (α and β) demonstrated comparable rescue effects, data was combined for all subsequent analyses **(Supplementary Fig. 2D)**. Exogenously expressed PSD-95 localized to dendrites and spine heads effectively, as we observed complete rescue of PSD-95 density in mutant neurons **(Fig. 3C-D, Supplementary Fig. 2E)**. To determine whether restoration of PSD-95 levels translates into structural changes in neurons, we measured dendritic spine density and observed rescue of earlier characterized deficits **(Fig. 3E)**. To further assess whether synaptic protein changes associated with DLG4 haploinsufficiency are affected by AAV9 gene therapy, we immunostained 30 DIV neurons for GluA2, AMPA receptor subunit 2, **(Supplementary Fig. 2F-H)**. Mutant neurons exhibited reduced GluA2 puncta density relative to wildtype, while AAV9 treatment restored GluA2 density, supporting recovery of PSD-95-mediated post-synaptic compartments.

### AAV9-hSynapsin-hDLG4 treatment rescues single-neuron and network functional phenotypes in a variant-dependent manner

Having established that AAV9-hSynI-hDLG4 restores PSD-95 levels and associated structural deficits, we next asked whether the electrophysiological signatures characterized above are similarly affected. To this end, we analyzed AAV9-hSynI-hDLG4 (α and β) infected cultures at 30 DIV, adding 31 recordings collected from two independent experiments comprising 2,453 units (HF AAV9, n=9 recordings/589 units; T654I AAV9, n=7/953; CH1371 AAV9, n=8/572; CH4053 AAV9, n=7/339), for a total of 88 recordings and 8027 spike-sorted units.

To examine the extent to which AAV9-treated cultures shifted toward the wildtype state, we projected AAV9-treated cultures onto support vector machine classifiers trained on mutant and wildtype cultures using single-cell, network-level and combined features, evaluating the percentage of recordings classified as “WT”, at 30 DIV. We observed a variable effect of treatment across feature groups, most notably in single-cell feature groups, as they separated cultures well previously (Fig. 2B), and across mutants **(Fig. 4A)**. This prompted us to explore the magnitude of the treatment effect, taking into consideration mutant vs WT separability, irrespective of classification outcome. To achieve this, we quantified the signed distance from the wildtype decision boundary and compared the performance of random forest classifiers using balanced accuracy **(Fig. 4B, Supplementary fig. 3A-C).** Firstly, classifier performance, quantified here as area under the precision-recall curve (PR-AUC), reproduced the feature dependence observed in our initial characterization, with the missense mutant lines separated principally by single-cell features and the lossof-function mutant lines by network-level features. Importantly, treatment shifted cultures toward wildtype in all four lines using combined features, particularly in the missense mutation (HF, T654I) lines **(Fig. 4B).**

**Figure 4:**
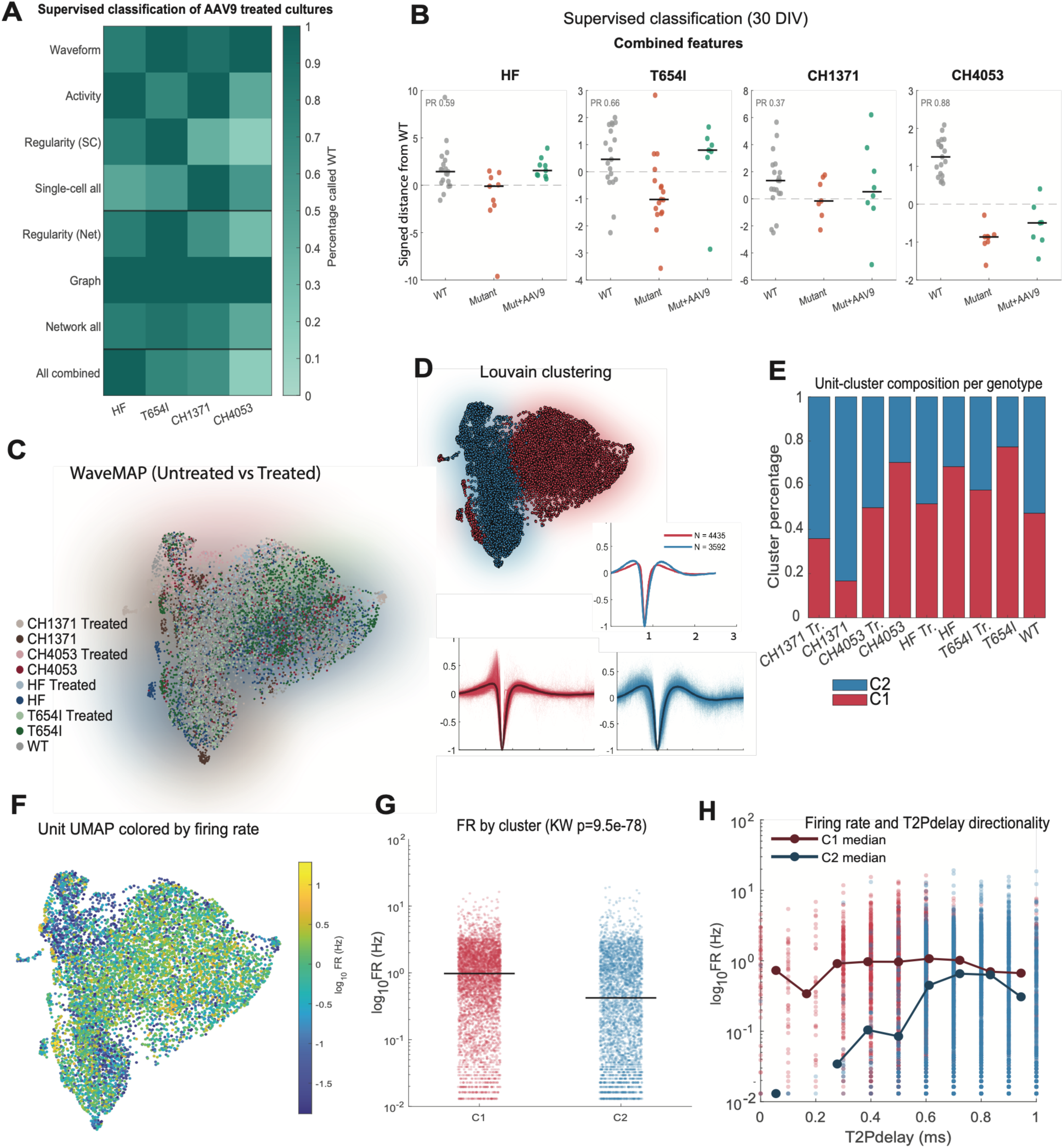
AAV9-mediated DLG4 restoration rescues single-neuron and network functional phenotypes in a variant-dependent manner. **(A)** Percentage of AAV9-treated recordings classified as WT per feature group and mutant, summarizing rescue across features. **(B)** Per-recording signed distance from the WT decision boundary using combined features for WT, untreated mutant, and AAV9-treated mutant, per mutant line. Precision- Recall AUC (mutant detection) indicated. Line in bars represents the median. **(C)** Single-cell waveform analysis (WaveMAP) embedding of per-unit waveforms colored by genotype and treatment. **(D)** Louvain clustering on WaveMAP from (C) and mean cluster waveforms identified from louvain clusters (Cluster 1: red, Cluster 2: blue). **(E)** Unit-cluster composition per group. **(F)** WaveMAP in (C) colored by mean firing rate per unit. **(G)** Average firing rate per cluster, Kruskal-Wallis, *p <0.0*5). **(H)** All units from WaveMAP in (C) plotted based on average firing rate and T2P delay values, colored by cluster origin.

To resolve these shifts at the level of individual neurons, we performed WaveMAP embedding of extracellular reference waveforms from all treated and untreated units followed by Louvain community detection, which resolved two waveform clusters **(Fig. 4C-D)**. Similarly to our earlier findings, the missense and nonsense lines were overrepresented in C1, the cluster comprising narrow-waveform units, while the frameshift line was underrepresented **(Fig. 4E).** Importantly, treatment moved all four lines toward the wildtype proportion, indicating that restoring *DLG4* normalizes the distribution of action potential waveform phenotypes, irrespective of the direction of the initial deficit. To confirm that these compositional shifts are not an artifact of pooling across genotypes, we performed recording-level and WaveMAP embedding separately for each loss-of-function line against its treated counterpart and wildtype **(Supplementary Fig. 3D–G)**, and for the missense mutation lines **(Supplementary Fig. 4A-D),** with WaveMAP analysis reproducing the shifts observed in the pooled embedding. These results motivated us to further explore how the signature waveforms translate into activity-level changes. Hence, we mapped firing rates onto the WaveMAP embedding of untreated and treated mutant and wildtype units **(Fig. 4F),** which revealed enrichment of higher firing units in genotypes overrepresented in C1, with significantly higher average firing rates than C2 **(Fig. 4G)** (p-val= 9.5e-78, Kruskal-Wallis). Moreover, this directionality was also observed at the unit level, as C1 units were concentrated at short trough-to-peak delays and fired at higher rates than C2 units, which occupied longer delays, indicating that the narrow-waveform and high-firing phenotypes co-segregate **(Fig. 4H).**

### Isolating the impact of the missense variant: Mutation-specific electrophysiological dysfunction

Having established variant-specific rescue effects across the mutant lines, we next sought to determine how well the missense variants phenocopied DLG4 haploinsufficiency. We first examined PSD-95 abundance in developing wildtype and patient-derived HF neurons by immunoblotting at days 9, 14, and 21 (**Fig. 5A).** Wild-type neurons showed progressive accumulation of PSD-95 from day 14 onward, whereas HF neurons exhibited markedly reduced PSD-95 levels, with only partial accumulation by day 21.

**Figure 5:**
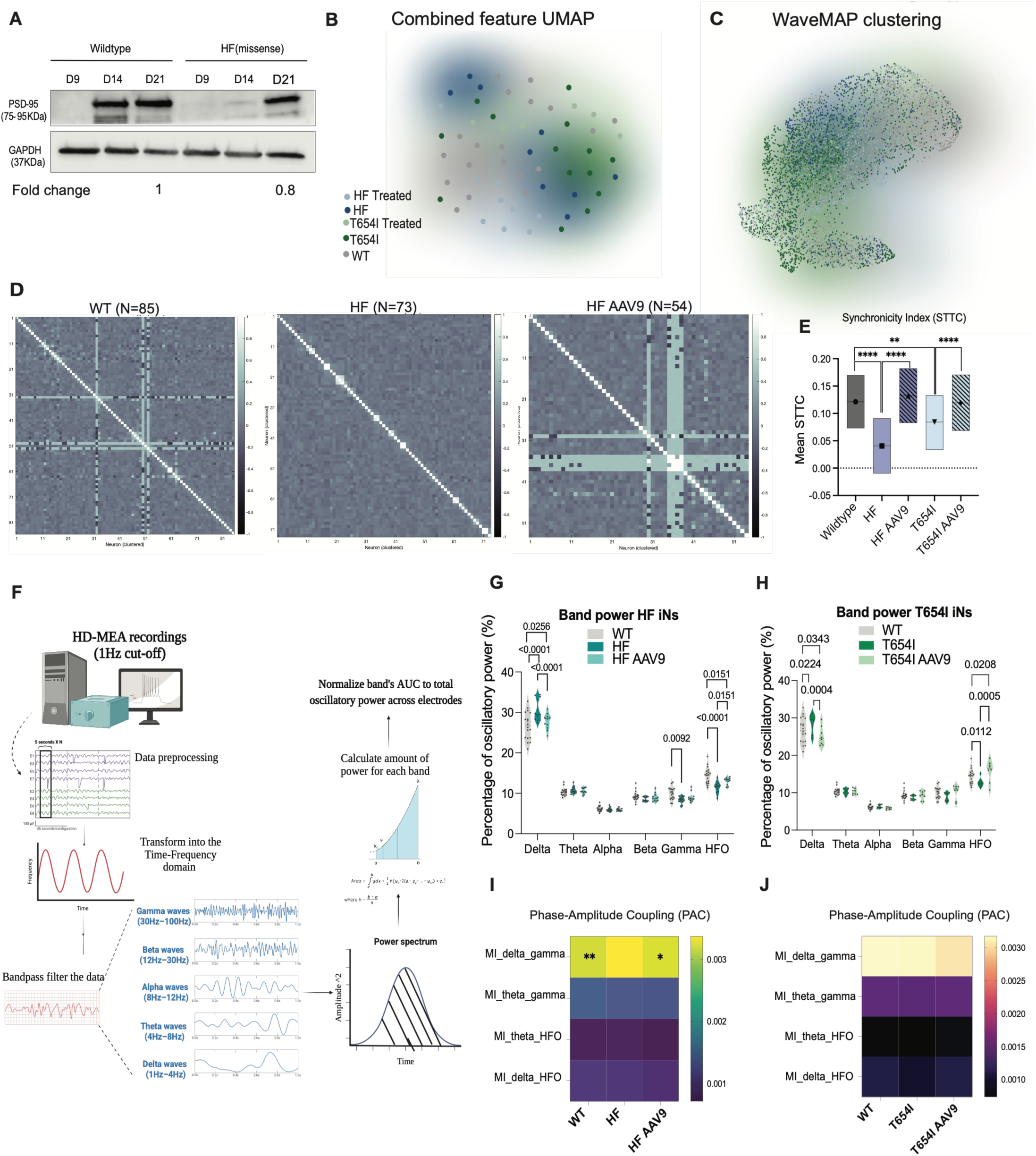
Isolating the impact of the missense variant: Mutation-specific electrophysiological dysfunction. **(A)** Western blot of PSD-95 in wild-type versus HF patient neurons across maturation. **(B)** Unsupervised clustering of mutant and treated cultures at the recording level across all features, at 30DIV. **(C)** WaveMAP embedding of per-unit waveforms by genotype and treatment. **(D)** Representative clustered spike-time tiling coefficient (STTC) matrices showing pairwise synchrony between spike-sorted neurons within individual recordings (N= neurons/well indicated). **(E)** Mean pairwise STTC values per group, Δ t=50 ms. Estimated marginal means and upper and lower CI are presented, line at median. Statistical analysis was determined using linear-mixed effects model (R, lmer package). Pairwise comparisons were analyzed with Bonferroni correction. Data collected from 2 (AAV9 treated mutants) and 4 (mutant and wildtype) independent experiments. Total wells and units analyzed: WT (n = 34 wells; 1667 units) pooled from three wildtype lines, HF (n =10; 801 units), HF AAV9 (n=9; 1246 units), T654I (n = 14; 999 units), T654I AAV9 (n =7; 576 units). **(F)** Schematic of LFP extraction and oscillatory band decomposition and power-spectrum analysis from HD-MEA recordings. **(G)** Oscillatory band power across bands for WT, HF, HF AAV9. **(H)** Oscillatory band power across bands for WT, T654I, T654I AAV9. **(I)** Mean modulation index (MI) values measuring cross-frequency coupling for WT, HF, HF AAV9. (**J)** Mean modulation index (MI) values measuring cross-frequency coupling for WT, T654I, T654I AAV9. Data collected from two independent experiments and represents individual wells. Statistical significance determined using Two-way ANOVA, corrected for multiple comparisons by controlling the FDR using BKY.

To separate the effect of the mutation from that of the patient genetic background, we applied DeePhys classification and WaveMAP analyses to the patient-derived HF and CRISPR-edited T654I missense lines, together with their AAV9-treated counterparts and wildtype controls **(Fig. 5B–D).** Combined feature embedding at the recording level revealed close clustering of the two untreated missense lines, whereas treated cultures shifted toward the wildtype **(Fig. 5B).** Similarly, WaveMAP analysis followed by Louvain clustering identified two neuronal clusters that were differentially represented across untreated mutant, treated, and wildtype cultures (**Fig. 5C, Supplementary Fig. 4E)**. Consistent with these cluster level shifts, raster plots of units grouped by WaveMAP cluster showed dense and temporally uniform firing in untreated missense cultures, compared to wildtype and treated cultures (**Supplementary Fig. 4F).** This phenotype was also evident at the single neuron level, with both missense lines exhibiting significantly higher firing rates and shortened interspike intervals, rescued by AAV9 treatment **(Supplementary Fig. 4G).**

To determine whether increased neuron-level firing rate was accompanied by altered coordinated network activity, we quantified pairwise spike time synchrony using the spike-time tiling coefficient (STTC)^28^. STTC matrices revealed reduced temporal coupling in both untreated missense lines compared with wildtype controls, whereas AAV9-treated cultures more closely resembled the wildtype pattern **(Fig. 5D).** Consistent with the latter, mean pairwise STTC values were reduced in untreated missense mutants and rescued following AAV9-treatment **(Fig. 5E)**.

Given the generalized background slowing observed in the HF patient (Supplementary Fig. 1A), we next examined whether the missense variants altered population-level oscillatory activity. Human brain organoids have been shown to generate field potentials from collective transmembrane currents that phaselock to neuronal spiking^29^, while phase–amplitude coupling has been detected in 2D human neuronal cultures recorded using multiwell MEAs^30^. LFP components were therefore extracted from raw HD-MEA recordings acquired with a 1-Hz high-pass cutoff, low-pass filtered at 300 Hz, and analyzed using custom MATLAB scripts **(Fig. 5F).** Spectral analysis revealed a shift toward slow-wave and low-frequency activity in both missense lines, characterized by an increased relative contribution of delta-band power in HF and T654I cultures **(Fig. 5G,H),** with AAV9 treatment of PSD-95 reducing delta-band power toward wildtype levels in both lines. To determine whether the shift toward low-frequency activity was accompanied by altered cross-frequency interactions, we quantified phase–amplitude coupling (PAC) between delta- and theta-band phases and gamma- and high-frequency oscillation amplitudes using the Modulation Index (MI)^31^. Among the frequency pairs examined, HF cultures exhibited significantly increased delta–gamma PAC, which was reduced following AAV9 treatment **(Fig. 5I).** T654I cultures showed a similar pattern, although the differences did not reach statistical significance **(Fig. 5J).** These results underscore the efficacy of AAV9-hSyn-DLG4 restoration in rescuing variant-specific electrophysiological signatures in *DLG4* missense mutant neurons.

## Discussion

Here, we provide, to our knowledge, the first experimental characterization of pathogenic DLG4 variants in human neurons, establishing a cellular model of DLG4-related Synaptopathy. Across multiple unrelated and isogenic wildtype and mutant iPSC-derived excitatory neurons, three distinct SHINE variants converged on a reduction in PSD-95 protein abundance. This reduction occurred despite variant-specific differences in mutant transcript levels, suggesting that haploinsufficiency in DLG4 is associated with decreased PSD-95 abundance, whether through transcript loss, protein instability, or both. This shared molecular consequence of pathogenic *DLG4* variants is consistent with higher constraint for LOF and missense *DLG4* mutations in population-level data^32^, and with enrichment of largely protein-destabilizing missense mutations with haploinsufficient genes^33^.

Reduced PSD-95 abundance was accompanied by a shared decrease in mature spine density across the mutant lines, consistent with impaired spine maturation or stabilization relative to wildtype neurons. This finding is supported by longitudinal *in vivo* imaging showing that recruitment and maintenance of PSD-95 is associated with spine maturation and increased stability, whereas low synaptic PSD-95 is associated with a higher probability of spine elimination^34^. In support of the latter, Yusifov et al. also revealed that PSD-95 knockout neurons exhibit increased spine elimination during experience-dependent structural remodeling^35^, and acute PSD-95 knockdown impairs activity-dependent spine maturation and maintains elevated spine turnover^36^. On the contrary, PSD-95 overexpression has been shown to promote glutamatergic synapse maturation and increased spine number and size^37^.

Consistent with the reduction in mature spine density, *DLG4* mutant neurons also exhibited reduced GluA2 density along dendrites, which was rescued following AAV9-mediated DLG4 restoration. Mechanistically, PSD-95 regulates synaptic AMPAR abundance through its interaction with stargazin, an AMPAR auxiliary protein, promoting receptor recruitment and stabilization at postsynaptic sites^38,39^. Importantly, mice expressing PSD-95 with ligand-binding-deficient PDZ1 and PDZ2 domains exhibit reduced GluA1 and GluA2/3 abundance in hippocampal PSD fractions across development, supporting impaired synaptic accumulation of AMPARs^40^. Moreover, PSD-95 deficiency impairs maturation of silent synapses, resulting in the persistence of synapses lacking functional AMPAR-mediated transmission^41^. Thus, reduced dendritic GluA2 density in *DLG4* mutant neurons may reflect impaired recruitment or stabilization of AMPARs at developing spines, contributing to fewer mature glutamatergic synapses.

These synaptic changes were paralleled by functional alterations. Unsupervised and supervised analyses of electrophysiological features of wild-type and mutant *DLG4* neurons revealed convergent phenotypes between nonsense and missense mutants, as well as variant-specific signatures in frameshift mutant neurons. Intriguingly, reference waveform-based analysis resolved the mutant cultures into two neuronal states defined by waveform shape, with narrow-waveform units showing higher firing rates and broadwaveform units showing lower firing rates. This waveform–firing relationship was differentially represented across *DLG4* genotypes and shifted toward wildtype following AAV9-treatment, suggesting that reduced PSD-95 abundance alters the balance between functionally distinct neuronal populations.

The observed shifts in waveform shape may reflect altered regulation of repolarizing conductances^42^. Prior studies provide a potential mechanistic link to PSD-95, which directly interacts with Kv1.4 containing Shaker-family of channel proteins and promotes their surface stabilization by limiting internalization ^13,43^. PSD-95 also binds the C-terminal VSAL motif of Kv4.2, with co-expression increasing channel surface abundance and clustering in heterologous cells^44^. Importantly, Kv4.2 and Kv4.3 mediated A-type currents contribute to rapid action-potential repolarization, and loss of either subunit prolongs action-potential duration in mature cortical pyramidal neurons. Notably, Kv1.4 deletion instead produces narrower action potentials through a compensatory enhancement of Kv4-mediated currents^45^. Thus, reduced PSD-95 may alter the balance and membrane organization of A-type potassium channel conductance, potentially contributing to the lower T2Pdelay observed in the narrow-waveform mutant state, while the broadwaveform enrichment in the frameshift mutation may reflect a distinct pattern of channel dysregulation or compensation.

Beyond altered A-type Kv conductances, extracellular waveform shape is also determined by the timing and spatial distribution of perisomatic transmembrane currents and by morphology-dependent filtering of these signals^46–48^. In this context, the selective reduction in ramification index in the frameshift line may contribute to its broad-waveform enrichment by altering dendritic current flow and extracellular filtering. This supports a shared mechanism underlying *DLG4* haploinsufficiency, that is influenced by phenotype-modifying factors, whether genetic, epigenetic, environmental, or developmental^49^ and contributes to the observed lack of genotype-phenotype correlations in *SHINE* syndrome.

Importantly, AAV9-hSynI-DLG4 restoration provides causal support for haploinsufficiency, as increasing PSD-95 abundance normalized synaptic markers and spine density and shifted electrophysiological signatures toward the wildtype state. The variant-dependent extent of rescue further suggests that reduced PSD-95 is a central pathogenic driver, while individual variants or genetic backgrounds impose additional constraints on neuronal maturation and circuit organization.

Given preservation of the mutant transcript, we examined the missense variant in greater depth and identified two main features. 1- High phenotypic similarity between the patient-derived HF and CRISPRedited T654I lines, evident by network and unit-level UMAP embeddings and multi-class supervised classification. Both lines also exhibited increased firing rates along with reduced pairwise STTC, indicating neuronal hyperactivity coupled to impaired temporal coordination, as reported in the autism model of TSC2 patient-derived neuronal culture^50^. 2-Recapitulation of patient-associated phenotypes. Both missense lines exhibited increased relative delta-band power, mirroring the generalized background slowing reported in the HF patient, with AAV9 treatment shifting this activity toward wildtype. Increased slow-frequency activity has also been reported in focal epilepsy, including enhanced delta and theta power over epileptic regions and increased scalp delta power associated with hippocampal interictal discharges^51,52^. HF neurons additionally revealed increased delta–gamma phase–amplitude coupling, which was reduced following AAV9 treatment, whereas T654I neurons showed a similar but non-significant trend. Abnormal PAC has been reported in epilepsy, autism spectrum disorder and genetic epileptic encephalopathy models, supporting its relevance to disrupted network organization^53–55^.

## Conclusion

The present study demonstrates the pathogenic consequences of *de novo* heterozygous DLG4 variants using patient-derived iPSC neurons. Molecular and electrophysiological analyses of mature excitatory neurons revealed convergent deficits across loss-of-function and missense mutant lines, including reduced PSD-95 abundance and associated structural and functional abnormalities. Importantly, AAV9-hSynI-mediated restoration of DLG4 rescued these abnormalities, supporting reduced PSD-95 availability as a shared pathogenic mechanism across variants. Together, these findings establish a human neuronal model of DLG4-related synaptopathy and support AAV9-hDLG4 restoration as a potential therapeutic gene therapy strategy.

## Materials and Methods

### Stem cell culture

Patient-derived induced pluripotent stem cell (iPSC) lines were obtained from established reprogramming sources through family-mediated access, table summary of iPSC lines included in the study present below. Stem cell lines were confirmed mycoplasma free and karyotyped normal, and maintenance was done on either feeder or feeder-free conditions. The former was performed using mouse embryonic fibroblast feeder layers seeded on 0.2% gelatin, and supplied with 20% KoSR primed media and 10 ng/ml bFGF, with daily media changes. For feeder-free culture, cells were plated on growth factor-reduced Matrigel and maintained in mTeSR1 medium with daily media changes. Cultures were routinely passaged at 70–80% confluency using TrypLE Express and supplemented with 10 µM ROCK inhibitor for 24 hours.

| iPSC line | Genetic background (NM_001365.5) | Source |
| --- | --- | --- |
| F1 | Wildtype | In-house generated |
| isoHF | Wildtype (isogenic HF line) | Hope for Harvey foundation |
| WT DLG4 | Wildtype (isogenic T654I line) | Hope for Harvey foundation |
| T654I | p.(Thr654Ile) (CRISPR-edited) | Hope for Harvey foundation |
| HF | p.(Thr654Ile) (patient-derived) | Hope for Harvey foundation |
| CH1371 | p.(Gly152Alafs*12) (patient-derived) | Children's Hospital of Eastern Ontario (DLG4 SHINE foundation) |
| CH4053 | p.(Arg352*) (patient-derived) | Children's Hospital of Eastern Ontario (DLG4 SHINE foundation) |

### Directed neuronal differentiation

iPSC-derived 2D neuronal cultures were derived through the previously established and characterized method of direct differentiation using NGN2 transcription factor overexpression (Zhang et al, 2013). Briefly, iPSC cultures were grown to 80% confluency prior to single cell dissociation and seeding in Matrigel (GFR)-coated 10cm plates, at 8e6-1.5e7 seeding density, in Neural differentiation medium (NDM), consisting of DMEM-F12, 1X N2 supplement, 1% Pen-strep, 1% NEAA, doxycycline (2ug/ml), BDNF (10 ng/ml), and NT-3 (10ng/ml). Differentiation was carried out for three full days with daily full media changes, after which cells were trypsinized and re-plated in neural maturation medium (NMM) consisting of base media supplemented with B27 with VA supplement (1X), Pen-strep (1%), Glutamax (1%), doxycycline (2ug/ml), mouse laminin (2ug/ml), BDNF (10mg/ml) and NT-3 (10ng/ml). Base media was BrainPhys medium for electrophysiological recordings, or Neurobasal medium. For the latter, final NMM was mixed with mouse astrocyte-conditioned NMM, in 1:3 ratio. Seeding cell densities are as follows: For immunofluorescence staining, 2e5 cells were seeded on 12 mm acid-washed glass coverslips, double-coated with 0.1% PEI and 10ug/ml mouse laminin. For biochemical assays, atleast 1e6 cells were seeded per well of 6-wellplate, and for HD-MEA recordings, 5e5 cells in 50 ul of media were plated covering the entire electrode array area. Cells were treated with 1 uM Ara-C and 10nM ROCKi for the first 48 hours only, with half media changes performed every three days for the remainder of the experiment.

### Immunocytochemical staining and analysis

NGN2-derived neurons, plated on glass coverslips, were washed with 1X PBS++ (with Ca and Mg) three times, fixed with ice-cold methanol for 20 minutes, then washed with 1X PBS++ three times. For brain cryosections, mice were anesthetized 2-3 weeks post injection, perfused with PBS, and fixed overnight with 4% PFA, then transferred to 30% sucrose for 24 hours, before embedding in OCT media for cryosectioning. For all tissues, permeabilization was done with 0.1% Triton X-100 in PBS (PBT) to enhance nuclear antibody entry, followed by blocking with 1% BSA and 5% goat serum in PBT for 1 hour at room temperature, and overnight primary antibody incubation at 4 degrees. Antibodies used: antiPSD5 (rabbit polyclonal, 1:250 dilution, #APZ-009), anti-PSD-95 (mouse monoclonal, 1:80, sc-32290), anti-GLUR2 (rabbit, 1:400, #AGC-005) anti-MAP2 (chicken polyclonal, 1:2000, ab92434).

Images were acquired using the Zeiss LSM-980 Airyscan, SR setting at 63X oil objective using 0.15-1 um intervals depending on culture density and preparation. Quantification of PSD-95 puncta number and volume was performed using synPAnal software^56^, on merged Z-stacks, automatically detecting thresholded puncta on highlighted 10 µm dendritic regions labeled with MAP2, whereas spine detection was manually performed on the same 10 um dendritic regions. Sholl analysis was performed through SNT tool, FIJI software^57^, using reconstructions of MAP2-labeled neurons generated through Vaa3D software^58^.

### Protein preparation and western blot analysis

For neurons, a minimum of 1.5e^6^ cells, were retrieved after trypsinization, then PBS washed before centrifuging at 1500 rpm for 10 minutes, then snap-frozen using liquid nitrogen. For mouse brains and organoids, tissues were lysed after snap-freezing. Pellets were subsequently lysed in commercially bought RIPA buffer supplemented with protease and phosphatase inhibitor cocktails on ice for 30 mins. Lysates were very briefly sonicated 2X (30 sec on, 45 sec off) prior to centrifugation at speed of 14000 rpm, at 4°C for 25 mins. Supernatant containing the proteins was retrieved and quantity was determined with Bradford assay. Samples (30-40µg) were denatured with 1X Laemmli sample buffer and boiled at 90 °C for 10 minutes. Samples were run on SDS-PAGE denaturing conditions on 10% TRIS-based separation gel, transferred to 0.2 um nitrocellulose membranes using the BioRad’s semi-dry transfer system. Membranes were blocked in 5% skim milk for 45 minutes prior to overnight antibody incubation at 4 °C under shaking conditions. Antibodies: PSD95 (mouse monoclonal, 1:500, sc-32290), GAPDH (mouse monoclonal, 1:1000, CB1001), GFP (goat polyclonal, 1:500, ab6673).

### RNA extraction and droplet-based digital PCR analysis (ddPCR)

RNA extraction was carried out using TRIZOL reagent, following manufacturer protocol. Briefly, cells were collected, washed with PBS, pelleted and snap frozen. For extraction, TRIZOL was added to frozen cell pellet and homogenized completely using a 21-gauge needle. Chloroform was added and mixed efficiently for phase separation of genomic material from organic protein layer. Clear upper aqueousphase was retrieved, and RNA was precipitated with isopropanol and glycogen and cleaned with multiple washes of 75% cold ethanol. RNA pellet was re-suspended in DEPC-treated RNase-free water, and 100500 ng was used for cDNA synthesis using the QScript cDNA synthesis kit. For digital PCR set-up, mutation-specific primer and probe sequences were generated using Primer3 website and SnapGene software. Primer sequences were mapped to span the exon-junction nearest to the mutation, with amplicon sizes between 75-150 bp, GC content 50-60% and Tm range of 55-60°. Wildtype (HEX) and mutant (FAM) probe sequences were mapped to center each mutation with a Tm=60°, with the addition of an MGB moiety. Each mutation run included a “no template” negative control well and two wildtype lines, along with patient neurons, all run in duplicate wells, on a QX200 ddPCR instrument (BioRad). Analysis was performed using BioRad Quantasoft software license.

### Oligo name Oligo type 5’ Dye Oligo sequence 3’ Modification

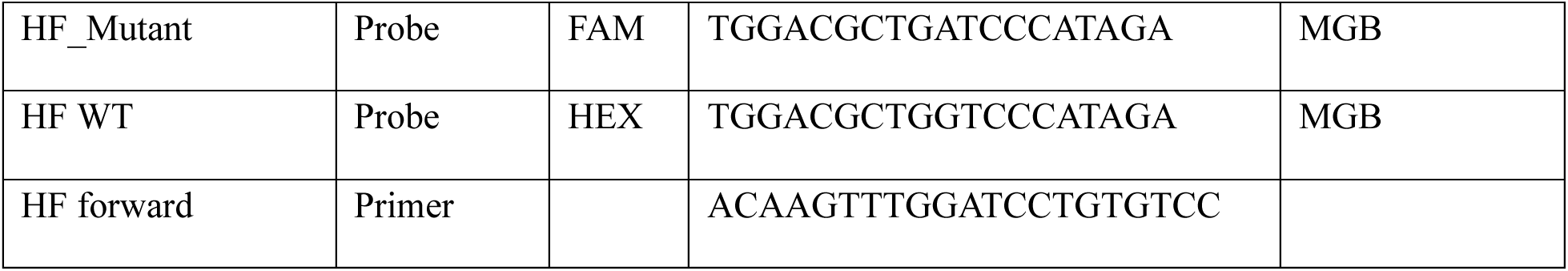
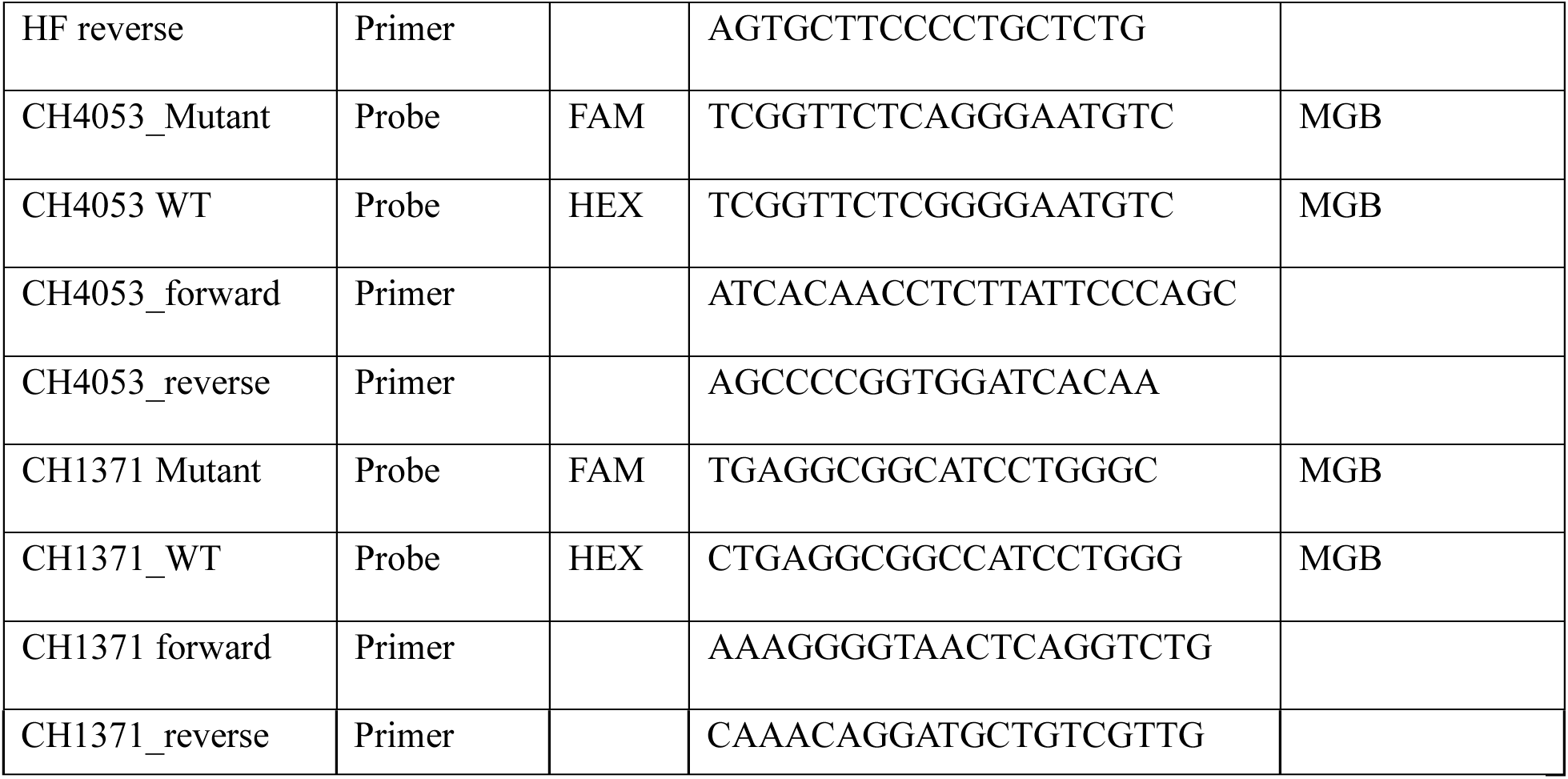

### Electrophysiological recordings

Extracellular recordings from neurons were acquired using MaxWell Biosystems MaxTwo recordings system, at a resolution of 26,400 electrodes per recording area, in physiological conditions of 5% CO2, and 37° temperature. Prior to each recording, MaxTwo well plates were left to acclimate in system incubator for 10 minutes. At every recording session, an activity scan of the full array configuration was performed, to identify active electrodes. A network assay based on electrodes with the highest spike amplitude was then performed, optimizing for spike sorting using the “Neuronal Units” configuration, for 5-10 minutes.

### Electrophysiological data analysis

For single neuron-level analyses, network assay raw data was exported into SpikeInterface^59^ python module, and spike sorted using Kilosort4 python package,^60^ using default parameters.

For analyses utilizing Deephys tool^26^, KS4 output data was loaded into MATLAB, and data was preprocessed using default parameters for feature extraction and subsequent single-cell and network feature analyses, excluding units with mean firing rates ≤0.01 Hz, as previously described^61^ and units exceeding the waveform-noise threshold (qc_noise = 8). Data was batch normalized for across-mutation UMAP embeddings. To account for imbalance in datasets, RF and SVM classifiers were class-weighted; RF using (Prior=’uniform’) and the SVM standardized using (Standardize=true).

Single unit firing rates, interspike intervals, and mean pairwise STTC values were computed using SpikenBurst tool^62^. KS4 output was QC’d and SpikenBurst GUI was set-up and analyze spike-sorted units, and default parameters were used. For synchrony analysis, STTC was used with 50 ms time window. Data was exported into excel and analyzed manually, using the mean unit-level metrics. STTC plots were generated through MATLAB, using the per-well STTC matrices from SpikenBurst. All single-unit data were analyzed using linear mixed-effects model to account for unit number without pseudoreplication, fitted by restricted maximum likelihood (REML) in R (lme4 with lmerTest). Data were log-transformed when residuals indicated a lognormal distribution. Genotype (treatment) was modeled as a fixed effect, with random intercepts for wells nested within batches to account for hierarchical, unbalanced, experimental structure (neurons nested within wells within batches). The model formula was: Data ∼ treatment + (1 | batch) + (1 | batch:well).

Degrees of freedom and p-values for fixed effects were calculated using Satterthwaite’s approximation. Estimated marginal means were calculated using estimated marginal means (EMmeans) and are presented with 95% confidence intervals. Pairwise comparisons were Bonferroni-corrected.

To examine extracellular local field potential (LFP) activity, raw data from HD-MEA recordings was exported into MATLAB and analyzed using custom-written scripts. Relative power was calculated for each frequency band, and phase–amplitude coupling was quantified using the modulation index based on Kullback–Leibler divergence.

### Statistical Analysis

Statistical analyses were performed using GraphPad Prism v.10. Data distribution was assessed within each group using the Shapiro–Wilk test. Normally distributed data were analyzed using one-way or two-way ANOVA, as appropriate, whereas non-normally distributed data were analyzed using the Kruskal–Wallis test. Multiple comparisons were performed using the post-hoc tests specified in the corresponding figure legends. Statistical significance was defined as P < 0.05. Data presentation, sample size, definition of n, and number of independent experiments are provided in the corresponding figure legends. Statistical analysis of single-neuron electrophysiological data is described separately in the electrophysiological data analysis section.

## Data Availability

Custom written MATLAB codes generated from this study can be supplied upon request from the corresponding author.

## Acknowledgements

We thank the Hope for Harvey Foundation for providing the missense mutant and isogenic iPSC lines. We would also like to extend our thanks to Dr. Alex MacKenzie and the CHEO Research Institute for providing the nonsense and frameshift patient iPS lines. We thank Ms. Lana Saffouri for technical help. Importantly, we are extremely grateful for the entire SHINE patient community for their perseverance and support.

This study was supported in part by the Hope for Harvey Foundation and the SHINE Syndrome Foundation.

## Author contributions

D.A. conceived and performed the majority of the experiments, analyzed and interpreted the data, prepared the figures, and wrote the original draft of the manuscript. M.O. performed the AAV9 injections in mice. R.I.A. conceived and supervised the study, secured funding, interpreted the data, and reviewed and edited the manuscript. All authors reviewed, revised, and approved the final manuscript.

