## Supplemental Figures for "Modeling and targeting haploinsufficiency in *SHINE* syndrome"

(Supplementary figures 1-4)

**A**

| Cell line | Mutation | Protein change | Exon | Effect | Protein domain | Gender | Seizure type, age of onset | ID/GDD |
|---|---|---|---|---|---|---|---|---|
| HF | c.1961C>T | p.(Thr654Ile) | 19 | Missense | GKLD | Male | Increased slow-wave, seizures later in age | Severe ID, GDD |
| CH4053 | c.1054C>T | p.(Arg352*) | 11 | Nonesense | PDZ3 | Male | Focal seizures, 10 years | Severe ID, GDD |
| CH1371 | c.455delG | p.(Gly152Alafs*12) | 7 | frameshift | PDZ1 | Male | Focal, focal-bilateral, tonic, 9 years | Severe ID, GDD |

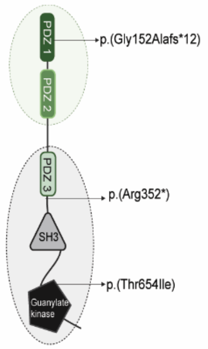


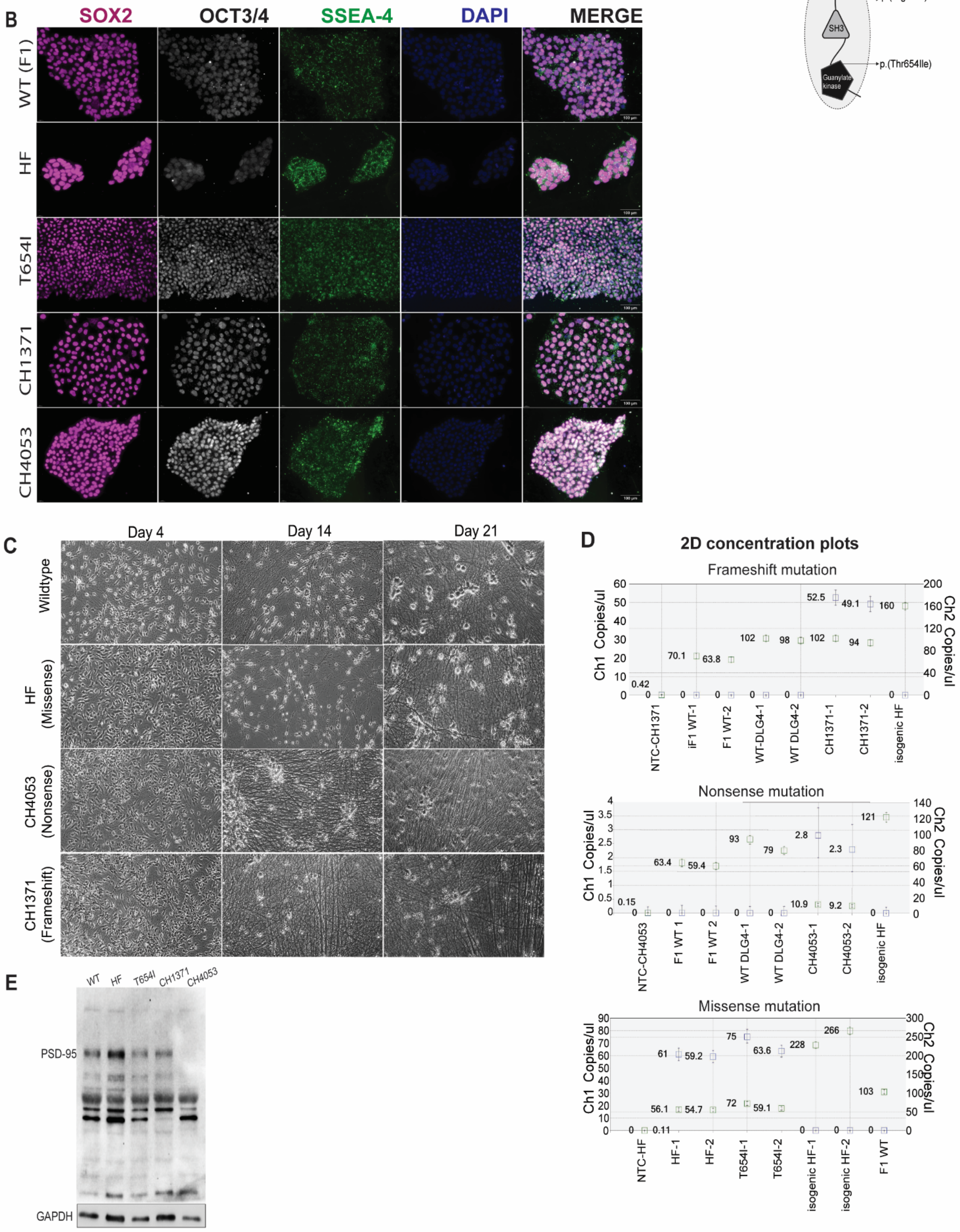

**Supplementary Fig.1:** Generation and neuronal differentiation of patient-derived hiPSC models of DLG4-related synaptopathy. **(A)** Left: Genetic and clinical features of the three patient lines carrying distinct *DLG4* variants. Right: Schematic of PSD-95 domains harboring the mutations (PDZ1–3, SH3, GK) with variant positions. **(B)** Immunofluorescent staining of pluripotency markers in F1 wildtype, and DLG4 mutant iPSCs. Scale bar= 100 um **(C)** Representative phase-contrast images of control and patient cultures across maturation of NGN2 neurons. Objective: 20X. **(D)** 2D concentration plots of mutant (channel 1), and wildtype (channel 2) transcripts of each mutant primer run. **(E)** Immunoblot of 30 DIV NGN2 neurons, showing full-length and degradation products of PSD-95.

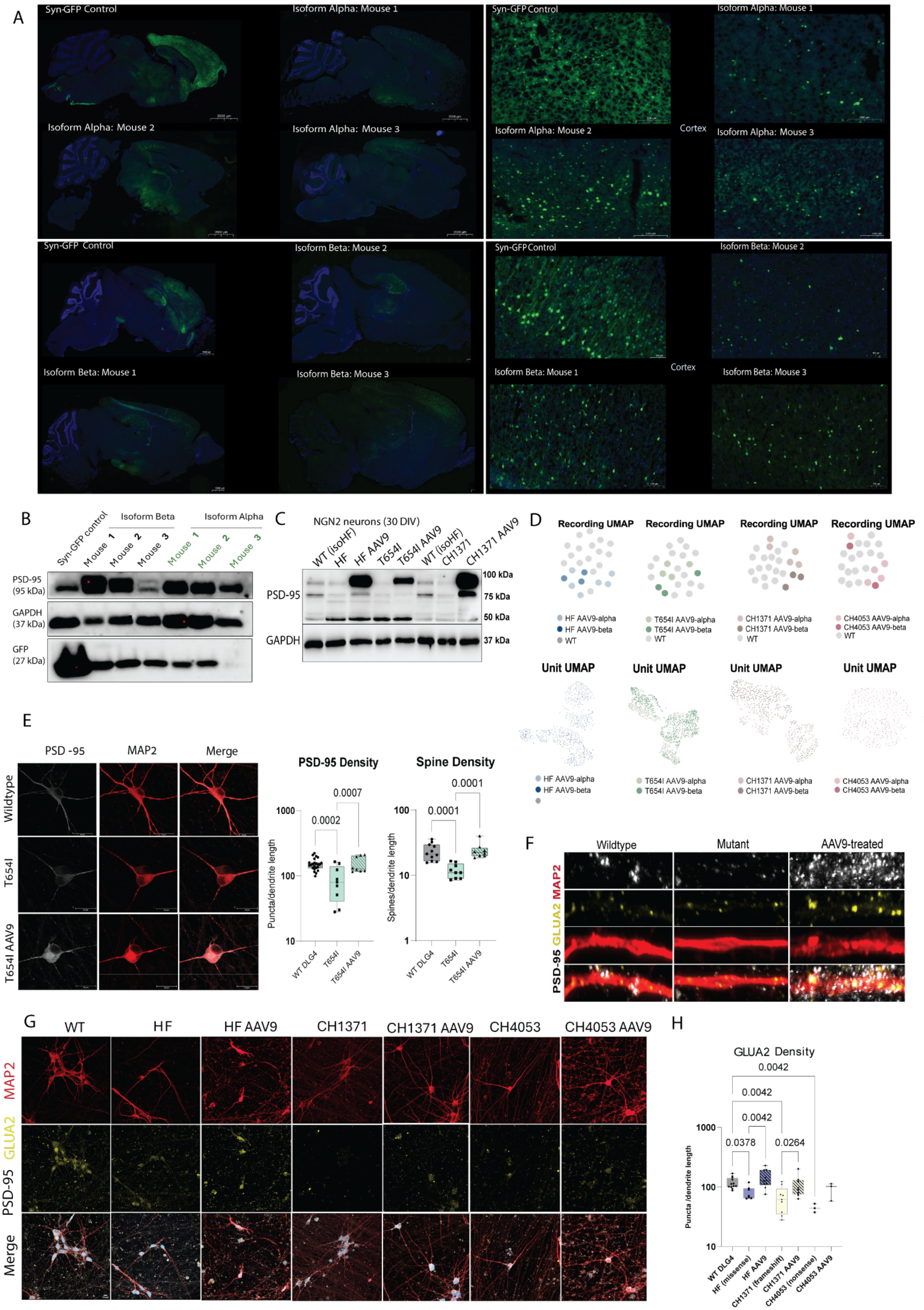

A
Syn-GFP Control
Isoform Alpha: Mouse 1
Isoform Alpha: Mouse 2
Isoform Alpha: Mouse 3
Cortex
Syn-GFP Control
Isoform Beta: Mouse 2
Isoform Beta: Mouse 1
Isoform Beta: Mouse 3
Cortex
B
Syn-GFP control
Isoform Beta
Mouse 1
Mouse 2
Mouse 3
Isoform Alpha
Mouse 1
Mouse 2
Mouse 3
PSD-95 (95 kDa)
GAPDH (37 kDa)
GFP (27 kDa)
C
NGN2 neurons (30 DIV)
WT (isoHF)
HF
HF AAV9
T654I
T654I AAV9
WT (isoHF)
CH1371
CH1371 AAV9
PSD-95
GAPDH
100 kDa
75 kDa
50 kDa
37 kDa
D
Recording UMAP
HF AAV9-alpha
HF AAV9-beta
WT
T654I AAV9-alpha
T654I AAV9-beta
CH1371 AAV9-alpha
CH1371 AAV9-beta
CH4053 AAV9-alpha
CH4053 AAV9-beta
Unit UMAP
E
PSD -95
MAP2
Merge
Wildtype
T654I
T654I AAV9
PSD-95 Density
Spine Density
Puncta/dendrite length
Spines/dendrite length
0.0002
0.0007
0.0001
0.0001
WT DLG4
T654I
T654I AAV9
F
Wildtype
Mutant
AAV9-treated
PSD-95 GLUA2 MAP2
G
WT
HF
HF AAV9
CH1371
CH1371 AAV9
CH4053
CH4053 AAV9
MAP2
PSD-95 GLUA2
Merge
H
GLUA2 Density
0.0042
0.0042
0.0042
0.0378
0.0264
Puncta /dendrite length
WT DLG4
HF (missense)
HF AAV9
CH1371 (frameshift)
CH1371 AAV9
CH4053 (nonsense)
CH4053 AAV9

**Supplementary Fig. 2: Validation of AAV9-hSynI-DLG4 expression and rescue of PSD-95-associated synaptic deficits. (A)** Representative sagittal sections and cortex regions from P30 mice following intracerebroventricular injection of AAV9-hSynI-GFP control or AAV9-hSynI-DLG4 expressing the human α-or β isoform. GFP is shown in green and nuclei are counterstained with DAPI. Three independently injected mice are shown for each DLG4 isoform. **(B)** Immunoblot analysis of PSD-95 and GFP expression in cortex of injected mice. **(C)** Immunoblot analysis of PSD-95 expression in 30 DIV untreated and AAV9-hSynI-DLG4(α)-eGFP treated NGN2 neurons. **(D) UMAP embeddings of AAV9-hSynI-hDLG4 (alpha) vs AAV9-hSynI-hDLG4 (beta) infected cultures showing no clear separation from wildtype recordings (top), and from each other at the unit level (bottom). (E)**Representative immunofluorescence images of wildtype, T654I and AAV9-treated T654I neurons stained for PSD-95 and MAP2. Scale bars= 20 um. Quantification of PSD-95 puncta and spine density along MAP2-positive dendrites in wildtype, T654I mutant, and AAV9-treated T654I. **(F)** High-magnification images of dendritic segments from wildtype, mutant and AAV9-treated neurons stained for PSD-95, GluA2 and MAP2. **(G)** Representative immunostaining of GluA2 and MAP2 in wildtype, HF, CH1371 and CH4053 neurons and their AAV9-treated counterparts. **(H)** Quantification of GluA2 puncta density along MAP2-positive dendrites. Statistical significance determined through One-way ANOVA, and adjusted for multiple comparisons by controlling the FDR using BKY. Datapoints represent technical and biological replicates across 2 independent experiments.

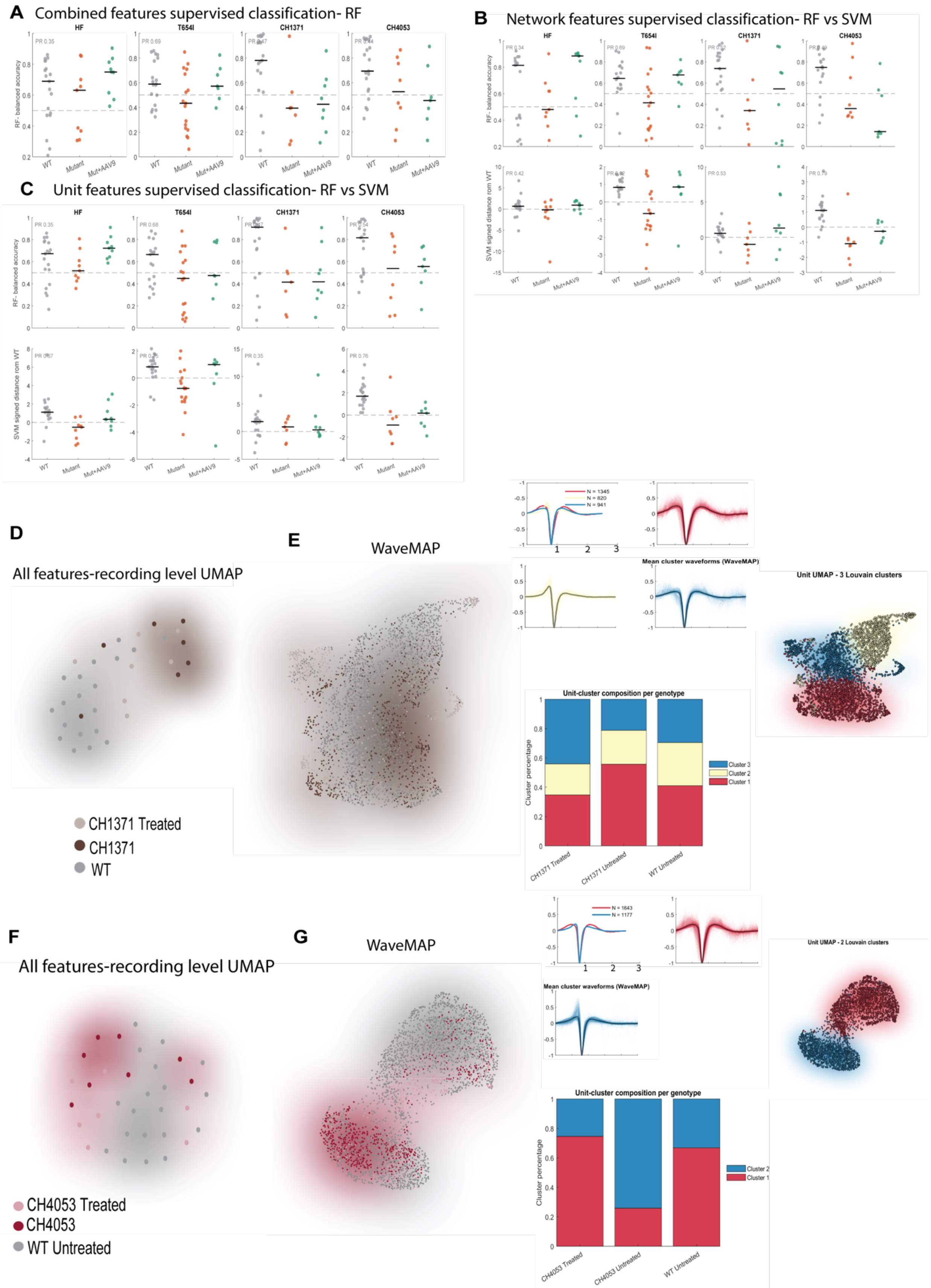
A Combined features supervised classification- RF
HF
T654I
CH1371
CH4053
RF- balanced accuracy
WT
Mutant
Mut+AAV9
B Network features supervised classification- RF vs SVM
SVM signed distance rom WT
C Unit features supervised classification- RF vs SVM
D
All features-recording level UMAP
CH1371 Treated
CH1371
WT
E
WaveMAP
Mean cluster waveforms (WaveMAP)
Unit UMAP - 3 Louvain clusters
Unit-cluster composition per genotype
Cluster percentage
Cluster 3
Cluster 2
Cluster 1
CH1371 Treated
CH1371 Untreated
WT Untreated
F
All features-recording level UMAP
CH4053 Treated
CH4053
WT Untreated
G
WaveMAP
Unit UMAP - 2 Louvain clusters
CH4053 Treated
CH4053 Untreated

Supplementary Fig. 3: **Feature-dependent classification and rescue of loss-of-function electrophysiological phenotypes**. **(A–C)** Supervised classification of wildtype, untreated mutant and AAV9-treated mutant recordings using (A) combined, (B) network-level and (C) single-unit feature groups. Random forest classifier performance is presented as balanced accuracy, with the dashed line indicating chance-level performance. Support vector machine classification is presented as the signed distance from the wildtype decision boundary, with zero indicating the decision boundary. Precision–recall area under the curve values is indicated. Each point represents one recording and black lines indicate median values. **(D)** Recording-level UMAP embedding of combined electrophysiological features from wildtype, untreated CH1371 and AAV9-treated CH1371 cultures. **(E)** WaveMAP embedding of CH1371, AAV9-treated CH1371 and wildtype units, followed by Louvain community detection. Mean waveform profiles and the relative composition of the three waveform clusters across groups. **(F)** Recording-level UMAP embedding of combined features from wildtype, untreated CH1371 and AAV9 treated CH1371 cultures. (G) WaveMAP embedding, mean waveform profiles and Louvain cluster composition of CH4053, AAV9-treated CH4053 and wildtype units.

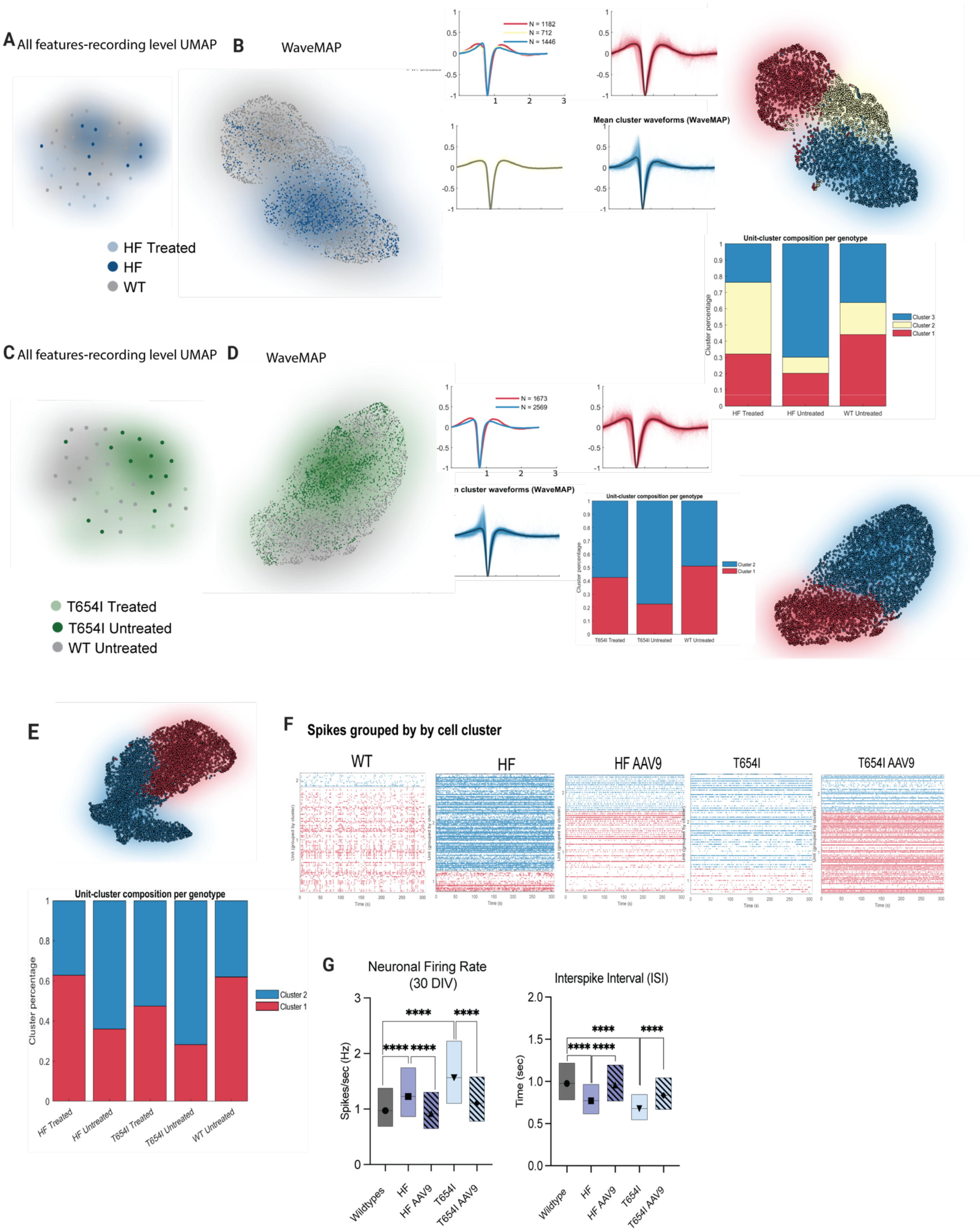

A All features-recording level UMAP
B
WaveMAP
HF Treated
HF
WT
N = 1182
N = 712
N = 1446
Mean cluster waveforms (WaveMAP)
Unit-cluster composition per genotype
Cluster percentage
Cluster 3
Cluster 2
Cluster 1
HF Treated
HF Untreated
WT Untreated
C All features-recording level UMAP
D
WaveMAP
N = 1673
N = 2569
T654I Treated
T654I Untreated
WT Untreated
E
Unit-cluster composition per genotype
Cluster percentage
Cluster 2
Cluster 1
HF Treated
HF Untreated
T654I Treated
T654I Untreated
WT Untreated
F
Spikes grouped by by cell cluster
WT
HF
HF AAV9
T654I
T654I AAV9
G
Neuronal Firing Rate
(30 DIV)
Spikes/sec (Hz)
Interspike Interval (ISI)
Time (sec)
****
Wildtypes
Wildtype
HF
HF AAV9
T654I
T654I AAV9

**Supplementary Fig. 4: Mutation-specific electrophysiological phenotypes and rescue in DLG4 missense mutant neurons. (A)** Recording-level UMAP embedding of combined electrophysiological features from wildtype, patient-derived HF and AAV9-treated HF cultures. **(B)** WaveMAP embedding of units from the same groups, followed by Louvain clustering. Mean waveform profiles and per-group cluster composition are shown. **(C)** Recording-level UMAP embedding of combined features from wildtype, CRISPR-edited T654I and AAV9-treated T654I cultures. **(D)** WaveMAP embedding, mean waveform profiles and Louvain cluster composition of T654I, AAV9-treated T654I and wildtype units. **(E)** Combined WaveMAP analysis of both missense models and their treated counterparts, showing the relative distribution of units between the two waveform clusters. **(F)** Representative spike raster plots from WaveMAP in (E) in representative recordings, showing proportion of units per well, colored by cluster-origin. **(G)** Estimated marginal means of neuronal firing rate and interspike interval at 30 DIV, analyzed using SpikenBurst tool. Statistical analysis was performed using linear-mixed effects model (R, lmer package). Pairwise comparisons were analyzed with Bonferroni correction. Data collected from 2 (AAV9 treated mutants) and 3-4 (mutant and wildtype) independent experiments.
